# Hybran: Hybrid Reference Transfer and ab initio Prokaryotic Genome Annotation

**DOI:** 10.1101/2022.11.09.515824

**Authors:** Afif Elghraoui, Deepika Gunasekaran, Sarah M. Radecke, Emma Bishop, Faramarz Valafar

## Abstract

De novo assembly has become commonplace for microbial organisms, increasing the demand for reliable genome annotation. Ab initio annotation is not an ideal approach for closely related strains due to suboptimal matching of the short or hypervariable genomic features that reference-based annotation transfer can overcome through identification of conserved synteny. At the same time, reference-based annotation methods leave gaps in the annotation where structural variations introduce unique sequence. We present Hybran, a hybrid reference-based and ab initio prokaryotic genome annotation pipeline that transfers features from a curated reference annotation and supplements unannotated regions with ab initio predictions. It builds on existing tools to create initial annotations using both approaches, then compares and resolves them to produce the hybrid annotation. With this pipeline, full advantage is taken of the community’s experimental efforts on reference strains to propagate as many known features as possible without sacrificing best-effort ab initio predictions for the remaining unannotated loci. Genome annotation performed in this way can facilitate comparative genomics and the investigation of evolutionary dynamics in microbial populations.

Hybran is freely available at https://lpcdrp.gitlab.io/hybran

## Introduction

Since the maturation of third-generation DNA sequencing technology(1), we have an abundance of completed microbial genomes. In particular, for sequencing projects that target populations of a single species to apply comparative genomics, having a set of genome annotations consistent across all members of a population would vastly simplify the task. Typically, for such organisms, a well-curated reference genome annotation exists, yet ab initio genome annotation methods are still used for new strains.

Although ab initio genome annotation methods ultimately query their detected open reading frames (ORFs) against reference databases, such searches can be difficult for genes that are hypervariable, short, or part of a gene family, such as the PE/PPE genes of the *Mycobacterium tuberculosis* complex(2) and the cytochromes of *Geobacter* species(3). Furthermore, a multitude of other genomic features such as transcription factor binding sites are not considered by some ab initio pipelines at all, mostly due to the unavailability of reliable generalized inference methods for them. Such features that have been experimentally identified for a reference strain can therefore only be propagated by reference-transfer via genome alignment.

Few tools for reference-based genome annotation exist. The only such tool that produces a complete result is the Comparative Annotation Toolkit (CAT)(4), a hybrid method, but its application is limited to eukaryotic genomes. The other remaining tools Liftoff(5) and the Rapid Annotation Transfer Tool (RATT)(6) are pure reference-based solutions, so cannot identify unique features in a strain. The incompleteness of their outputs has largely limited their application, so a pipeline to incorporate this approach while producing a more thorough annotation is needed for prokaryotic genomes. For this purpose, we present Hybran, a hybrid annotation pipeline leveraging conservation of synteny via RATT to maximize the benefits of highly-curated reference annotations and produce an ungapped result.

## Methods

### Overview of the Hybran Pipeline

Hybran first transfers annotations from one or more user-provided references. For best results, the reference annotations should be well-characterized and closely related to the genome sequence(s) to be annotated. Next, Hybran performs ab initio annotation, followed by merging the two results for each input genome into a draft hybrid annotation. Annotations discarded from either source are logged, together with the reason why. Finally, all reference and sample genes from these draft annotations are clustered together to potentially match functional annotations to hypothetical genes. Hybran has global settings for amino acid sequence identity and alignment coverage thresholds, used at various stages of the pipeline. The full pipeline is presented in Figure 1 and described in more detail in the following sections.

**Figure 1:**
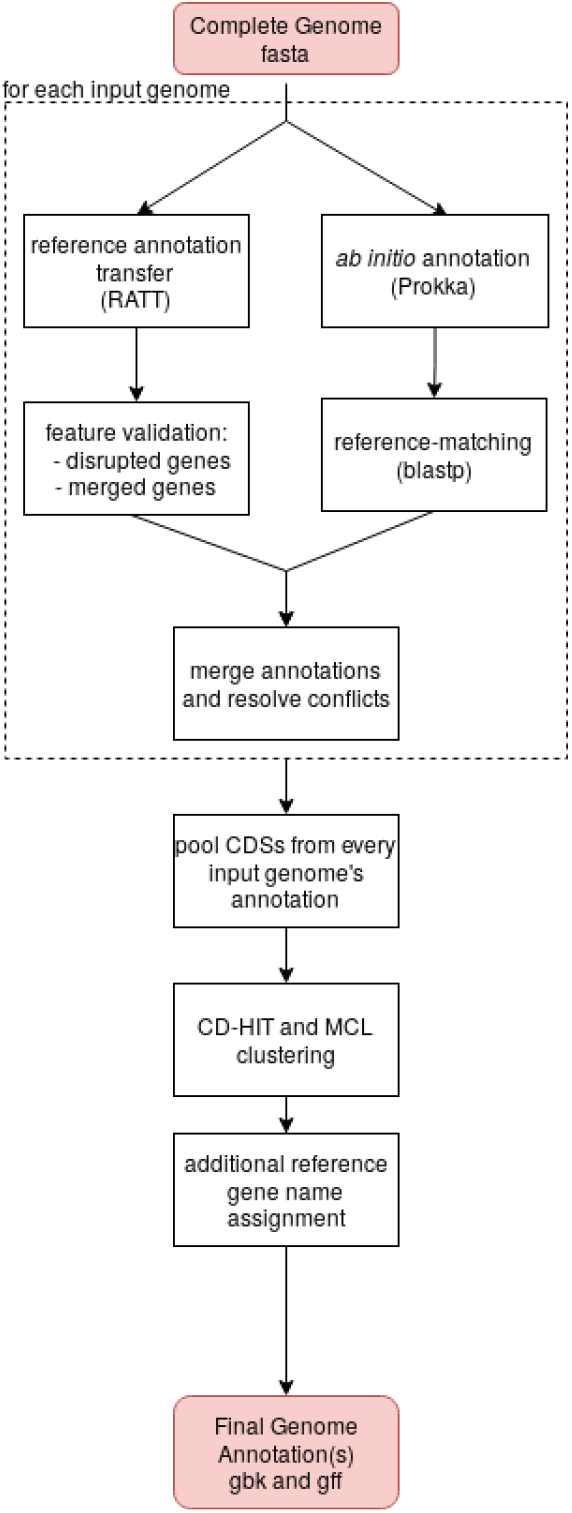
Overview of the Hybran Pipeline

### Preprocessing of Reference Annotations

Duplicated genes in reference annotations are either unnamed or distinguished from each other with number of letter suffixes. If one of these copies’ annotations cannot be transferred (for example, due to the ortholog being adjacent to a large insertion with respect to the reference), or if the sample has an additional copy of the gene than the reference does, these genes would not be consistently named. Furthermore, a defining feature of Hybran is that homologous genes in its output should be identifiable solely by looking for identical gene names.

To enhance this property, Hybran allows for an optional step of assigning a single gene name for each set of duplicated genes in the reference with the --dedupe-references flag. If this step is enabled, CD-HIT(7, 8) clustering is applied to reference amino acid sequences using 99% sequence identity, query and subject alignment coverage to collect sequences to be considered identical. All the genes in each cluster are then assigned a single generic name for use in the remainder of the pipeline. An updated reference annotation using these names and a list of duplicated reference genes are included among the output.

### Annotation Transfer

The first step in the annotation pipeline is the transfer of genes from the reference annotation(s) using RATT(6). RATT works by pairwise aligning the input genome to the reference sequences, identifying matching stretches of sequence, and then transferring the corresponding annotations while ensuring that gene start and stop codons remain accurate and correcting them in the case of mutations. The greatest advantage of this approach is that genomic features that are individually difficult to match can be matched by virtue of conserved synteny: features that may be small, polymorphic, or members of gene families.

RATT’s parameters include a transfer type, which represents the expected degree of similarity between the reference and the sample sets, and additional settings provided via an optional configuration file. These settings include a list of start and stop codons, splice sites, and Boolean settings regarding whether to attempt correction of splice sites and pseudogenes. All of these settings are directly configurable by the user through Hybran’s command-line interface, with the exception of the start and stop codon lists, as Hybran automatically detects the appropriate genetic code from the provided reference annotation. With these parameters, Hybran dynamically generates the configuration file for RATT prior to execution.

The output of RATT is validated and postprocessed to improve compliance with International Nucleotide Sequence Database Collaboration (INSDC) standards and to remove artifacts. In particular, when frameshift or nonsense mutations occur in a sample gene with respect to a reference, RATT currently annotates the gene using a compound interval to exclude internal stop codons, when it should rather use a single interval and apply the “pseudo” annotation tag to the record to indicate that something may be wrong with the gene.

Hybran then runs a BLASTP(9) alignment of each transferred gene to the reference copy and stores the alignment results (percent identity and query and subject coverages) primarily for later use in comparisons to the ab initio annotations during the annotation merging step. An option (--filter-ratt), however, exists should the user wish to enforce the global identity and coverage thresholds on the RATT-transferred annotations. But we do not recommend using this option since it largely negates the advantage conferred by synteny-based annotation transfer.

### Ab initio Annotation

In spite of high sequence conservation across strains, there may yet remain areas of the genome that have changed due to recombination, duplication, or spontaneous mutation. In these regions, RATT will not be able to transfer annotations since the sequence identity has altered significantly or the sequence does not exist in the reference, necessitating this ab initio component of Hybran, for which we currently use Prokka(10).

The pipeline runs Prokka with parameters --rfam and --rnammer. Prokka’s --coverage option is set to match Hybran’s alignment coverage threshold parameter, and Prokka’s --gcode flag is set to match the genetic code of the reference annotation as detected by Hybran. Most other settings of Prokka, such as the E-value cutoff, are exposed to the user and otherwise set to Prokka’s defaults.

Ab initio annotations are post-processed in two steps. First, they are compared to reference protein sequences and annotations are transferred for acceptable matches. Then, Hybran attempts coordinate correction by comparing the sequence context of reference orthologs to the reference genes themselves. These post-processing steps are detailed in the following subsections.

### Comparison to Reference Protein Sequences

While Prokka can be run in a way to prioritize, for functional assignment, BLASTP hits to user-provided annotation proteins with its --proteins option, we found Prokka’s matching criteria to be inadequate. In its BLASTP sequence alignments, Prokka accepts matches that meet its E-value and query coverage thresholds (--coverage). However, due to the lack of consideration of both subject (reference) and query alignment coverage, Prokka will annotate a coding sequence (CDS) with the same functional information as the subject sequence even though they were of widely disparate lengths and a strong E-value achieved because a shared sequence domain. Alternatively, a mutation in the sample would have introduced a premature stop codon, thereby truncating the protein with respect to the reference, which, although still warrants the functional assignment, the “pseudo” attribute should also be applied to the annotation.

Hybran compares CDSs to reference protein sequences using three metrics from BLASTP alignments: percent identity, query alignment coverage, and subject alignment coverage. In a set of criteria that are also applied to BLASTP alignments throughout the pipeline where Hybran is evaluating orthology, hits are evaluated according to the configured thresholds. If all three thresholds are met, the annotation is transferred. If percent identity and only one of the alignment coverage thresholds are met, the match is considered further. If the only deficient metric is query coverage, meaning that the query CDS is mostly similar to the reference CDS, except the query CDS is much longer, the annotation is transferred, as this may indicate that the reference CDS itself is truncated. If the only deficient metric is subject coverage, meaning again that the two sequences are mostly similar, but instead the query CDS is much shorter than the reference, then this is suggestive of a premature stop codon truncating the query CDS. In this case, the reference annotation is transferred, but with the addition of a “pseudo” tag to indicate a potential problem with this copy of the gene. In all other cases, no annotation is transferred and any gene name assigned by Prokka is removed, being assigned instead as a “gene_synonym” attribute, to prevent propagation of unreliable matches in the final clustering step.

### Correction of ab initio-predicted Gene Coordinates

For ORF-finding, Prokka utilizes the gene prediction algorithm Prodigal(11), an unsupervised machine learning algorithm that uses the input genome sequence to learn gene structure properties such as codon statistics, ribosome binding site (RBS) motif usage, start codon frequencies (only ATG, GTG and TTG) and predicts gene coordinates in this genome to identify candidate gene coordinates. Although utilization of genome-specific metrics such as codon statistics make Prodigal an unbiased tool for gene predictions, some implicit biases may occur due to restriction of start codons to ATG, GTG and TTG. If a rare codon is the start, Prodigal would continue searching until one of the three start codons was identified (11). This bias can be corroborated with the results of gene coordinate prediction of 62 experimentally verified genes in *Mycobacterium tuberculosis* H37Rv using Prodigal, where all 62 genes were identified with 58 having the correct start coordinate, indicating that the start site prediction by Prodigal has an accuracy of 93.6% for the set of 62 experimentally verified genes(11).

Because Prodigal was trained on a specific set of prokaryotic genomes and contains general rules about gene structure in prokaryotes(11), Hybran executes Prodigal on the provided reference genome and compares the output gene coordinates to those in the reference annotation. Gene coordinates with incorrect (according to the reference annotation) start positions are noted. When any of these genes are encountered in an input genome’s ab initio (Prodigal) annotation, the length of each gene was compared to the same gene in the reference. If the lengths were different, the upstream nucleotide sequence context (40bp) of this gene (the start was considered the same, relative start as that in the reference) was compared to the reference’s upstream context to check for any mutations that may have caused Prodigal to choose a different start coordinate. If a mutation in the upstream region of the gene was found, the ab initio gene coordinates were accepted; otherwise, the coordinates were recalibrated to match those of the reference. In *Mycobacterium tuberculosis* H37Rv, for example, we found Prodigal to have 88% concordance with the reference annotation: 480 open reading frames called by Prodigal had discordant start coordinates.

### Merging Transferred and Ab Initio Annotations

With the reference-transferred and ab initio annotations ready, the merging of the two can proceed. This step begins with the reference-transferred annotation as a starting point, excluding annotations for ribosomal RNAs (rRNAs) and transfer RNAs (tRNAs). We have found that rRNA annotations are not reliably transferred by RATT. As for tRNAs, they can be detected with high sensitivity by ab initio methods, so do not warrant as much the risk of RATT missing some calls due to lack of synteny in certain regions. The merging process is described in the following subsections. Any annotations rejected, from either source, are logged with justification.

#### Identifying Gene Fusion

The reference-transferred annotations are checked for signs of gene fusion. In RATT’s output, this manifests in two different ways:

1. Two different genes are annotated as having identical genomic coordinates. Two genes that were adjacent in the reference annotation are found to have sequence similarity to the sample. Through RATT’s start and stop codon correction, the two annotations end up with the same coordinates. If an ab initio annotation exists with these coordinates, its annotation is used here. Otherwise, the RATT annotation is maintained, but the gene name is removed from the annotation. In either case, Hybran adds a note to the annotation mentioning that the two genes have fused together.
2. Two different genes with the same stop position, but different start positions. If, with respect to the reference genome, the stop codon of a gene is lost due to mutation and the reading frame permits continuation through the rest of the gene, intergenic region, all the way to the stop codon of the next gene downstream, this case arises. The first gene is conjoined with the second, but the annotation of the second gene is intact. Hybran annotates the first gene with its original name, but adds a pseudo tag (a tag which, again, signifies a potential problem with the gene) and a note describing its fusion with the neighboring gene. The second gene’s annotation is maintained as is.

#### Discarding of Redundant ab initio Annotations

To reduce the number of overlapping features to consider when merging the annotations, Hybran makes a first pass identifying ab initio CDSs that are clearly redundant with the reference-transferred annotations. These are annotations that have start and stop positions each within 5 bp of each other and either have the same gene name (recall that ab initio CDSs are assigned reference gene names as part of our postprocessing) or have identical length. Such ab initio annotations are discarded at this stage.

#### Merging and Resolving of Overlapping Annotations

Ab initio features that either have no overlap with reference-transferred annotations or do, but are a different type (for example, RNA versus CDS) are automatically accepted for inclusion into the final annotation.

Before resolution of conflicting CDSs, the ab initio annotation is aligned with BLASTP to each reference-transferred annotation that it overlaps with (recall that the ab initio annotation was previously compared to the reference annotations by this point only). If there is a match according to Hybran’s criteria (see section *Comparison to Reference Protein Sequences*), the same gene name is assigned. A resolution between the two annotations has not been determined at this point, but the identity of the gene name ensures that Hybran later accepts only one of the two annotations.

The resolution proceeds for each overlapping pair of annotations according to the following inclusion criteria:

1. If, by this point, the two annotations do not share the same gene name (i.e., are not in the same open-reading frame) both are accepted.
2. If they do share the same name, the upstream sequence context of each gene (set as 40 bp upstream from the start position) is compared to the reference gene’s upstream sequence. This is to assess which has a more accurate start position according to the reference annotation.

a. If the RATT annotation’s upstream context matches exactly, the ab initio annotation is rejected.
b. If not, the start position accuracy is not necessarily called into question, as there may simply be a mutation in this region. Instead, each’s BLASTP hits to the reference are compared by using the absolute difference of the subject and query coverage for each. The annotation with the smaller difference is accepted. If these quantities are identical, the annotation that has the higher percent identity hit is accepted.

By this point, the merging of the reference-transferred and ab initio annotations is complete and postprocessing of the result begins.

#### Combining Putative Gene Fragments

Ab initio annotations may have called multiple adjacent ORFs that partially aligned to the same reference gene. In this case, these annotations would be adjacent on the same strand, share the same name, and each have a “pseudo” tag that would have been previously assigned through Hybran’s reference matching. Hybran makes a final pass through the final set of annotated genes and combines such entries into one annotation with the start position of the first putative fragment and the stop position of the last and the proper strand orientation.

### Clustering

To identify genes that have still not been assigned a gene name, the pipeline next clusters sequences first using CD-HIT(7, 8) and then with the Markov Cluster (MCL)(12, 13) algorithm. This leverages all hitherto identified orthologs and paralogs to potentially identify more. All genes from all (input and reference) genomes are used here, as this step runs once merged annotations have been produced for each input.

CD-HIT sorted a given set of sequences (here all CDSs present in the set of genomes) in descending length and the longest sequence was chosen as the representative sequence for the first cluster. Each sequence was then compared to this representative and considered part of the same cluster based on a 99% to 98% amino acid sequence identity and 100% coverage. If the query sequence did not meet these thresholds, it was considered the representative sequence of a new cluster. This process continued for all protein sequences in all the clinical isolates. The identity threshold was then reduced by 0.005% and the process began again based on the previous set of clusters, which created a cluster of clusters that was input into an all-against-all BLAST.

An all-against-all BLASTP alignment was implemented to create a matrix. The matrix represents a connection graph with sequences or CDSs as nodes and sequence identity as edges (greater than 95% identity and query and subject coverage). Weights were calculated by taking the logarithmic value (base 10) of the E-value from the BLASTP alignment between those two nodes or CDSs.

The matrix was then passed into MCL, which performs iterative rounds of matrix multiplication (expansion) and inflation (contraction) until there was no net change. To determine when there was no net change, the inflation parameter, which determines the granularity or tightness of the clusters, was set to 1.5 in our clustering step. The lower the inflation threshold the tighter the clusters (range of 1 to 10). This tighter threshold was chosen based on the higher-level genome conservation expected of the types of inputs this pipeline is designed for.

Based on the clusters created above, we isolated clusters that contained unnamed sequences, which resulted in three types of clusters: (1) clusters containing only unnamed sequences, (2) clusters with both commonly named sequences and unnamed sequences, and (3) clusters with differently named sequences and unnamed sequences.

Steps to assign names for each case are as follows:

1. *Clusters containing only unnamed sequences*: The representative sequence of this cluster was compared to all reference annotation CDSs using BLASTP and, again, Hybran’s criteria (see section *Comparison to Reference Protein Sequences*). If no hit is found, the cluster was assigned a new generic gene name, by default of the form ‘ORF#####’, where the ‘#’ indicates a digit, starting at ORF00001, assigned in order of the clusters output from MCL.
2. *Clusters with commonly named sequences and unnamed sequences*: All unnamed sequences in these types of clusters were assigned the gene name shared by the named members of the cluster.
3. *Clusters with differently named sequences and unnamed sequences*: Each unnamed sequence was compared to all named sequences in the cluster using BLASTP and Hybran’s global criteria to determine which named gene the unnamed sequence was most similar to. The unnamed sequences were annotated with a gene name based on the best hit to a named sequence. If the thresholds were not met, the gene is assigned a generic gene name.

The merged annotations for each input genome are then updated with the newly assigned gene names, finalizing the result.

### Evaluating Hybran

To evaluate Hybran’s performance, we compared its annotations to those produced by the NCBI Prokaryotic Genome Annotation Pipeline (PGAP). PGAP is a thorough and extensive pipeline, making use of multiple gene-finding algorithms and comprehensive collections of a variety of databases. We applied each tool to genomes of two different species: *Mycobacterium tuberculosis* H37Ra(14) and *Pseudomonas aeruginosa* PAK(15).

We ran PGAP with genome topology setting “circular” and setting the genus_species setting for each input accordingly. We ran Hybran with --genus and --species similarly set, as well as with the --dedupe-references flag (see *Preprocessing of Reference Annotations*). For the reference annotation of *M. tuberculosis*, we used that of strain H37Rv(16, 17). For *P. aeruginosa*, we used that of strain PA01(18, 19).

We compared the results of each tool using a modified version of Compare-annotations (https://github.com/rrwick/Compare-annotations). This script conducts a global alignment of gene sequences from one annotation source to those of the other and determines differences in annotation based on gene coordinates and, in its original form, the *product* fields of the annotations. Our modified version instead compares the gene names of each annotation, as the primary goal of Hybran is to maximize the extent to which genes from the reference strain, especially those still unnamed and uncharacterized (where the reference-specific locus tag serves as the gene name), can be identified in a genome.

## Results

We first provide a visual representation of the methods contributing to the final annotation output by Hybran (Fig. 2). In both cases, the majority of annotations are sourced from RATT, with ab initio annotations contributing to a few segments in each genome. The latter makes a greater contribution to the annotation of the PAK genome, as this has many more synteny-disrupting structural variations with respect to its reference genome than H37Ra.

**Figure 2.**
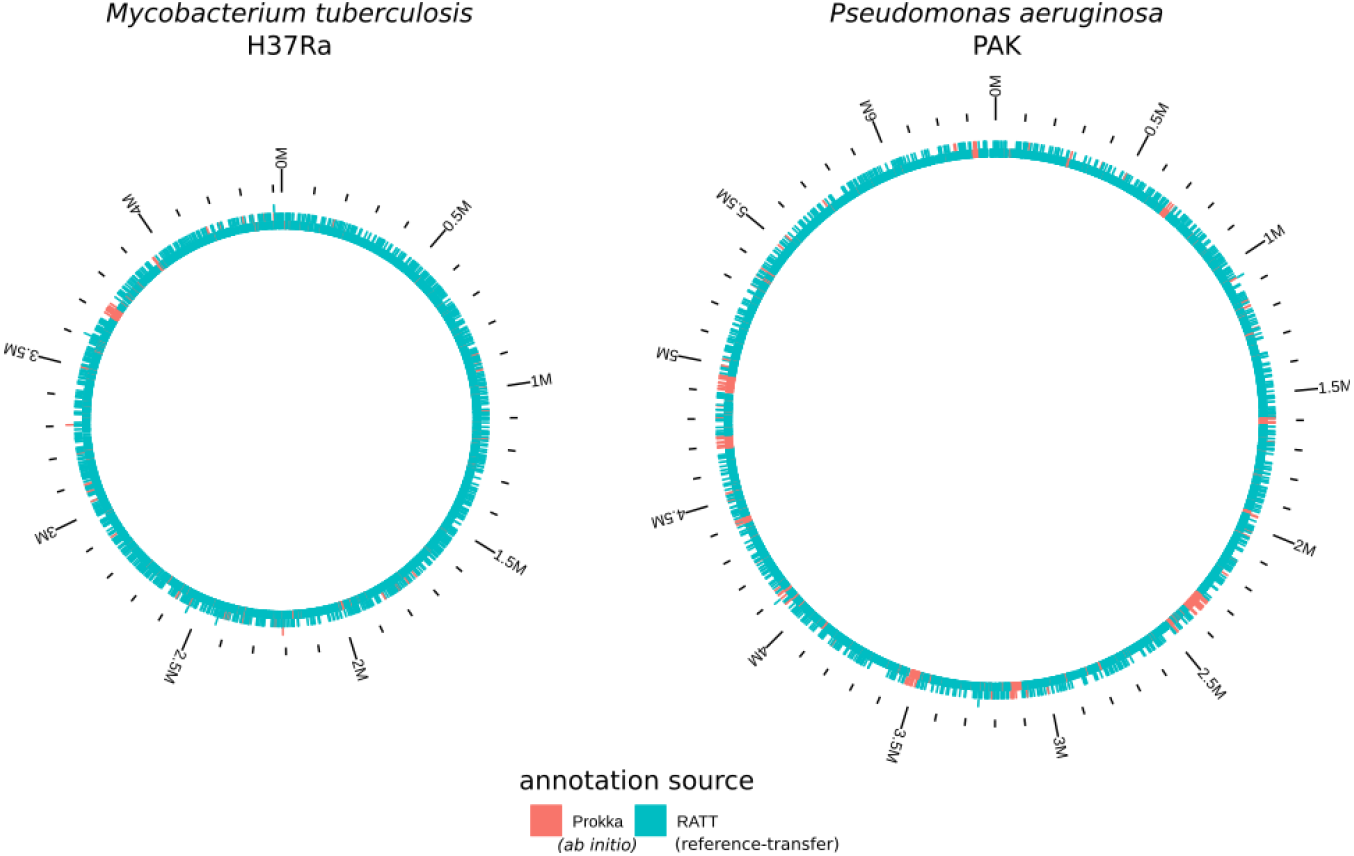
Sources of Final Hybran Annotations of *M. tuberculosis* H37Ra and *P. aeruginosa* PAK.

The results of the comparisons between Hybran and PGAP are shown in Table 1. For CDSs that both tools annotated, Hybran assigned reference gene names much more frequently. For PAK, 1787 out of the 5501 CDSs that both Hybran and PGAP annotated were only assigned names by one method: 5 by PGAP and the remaining 1782 by Hybran. For H37Ra, 612 out of 3476 common CDSs were named in only one pipeline, and all were by Hybran. We did not compare CDSs whose coordinates were partial matches between the two pipelines (197 CDSs for PAK and 458 for H37Ra).

**Table 1.**
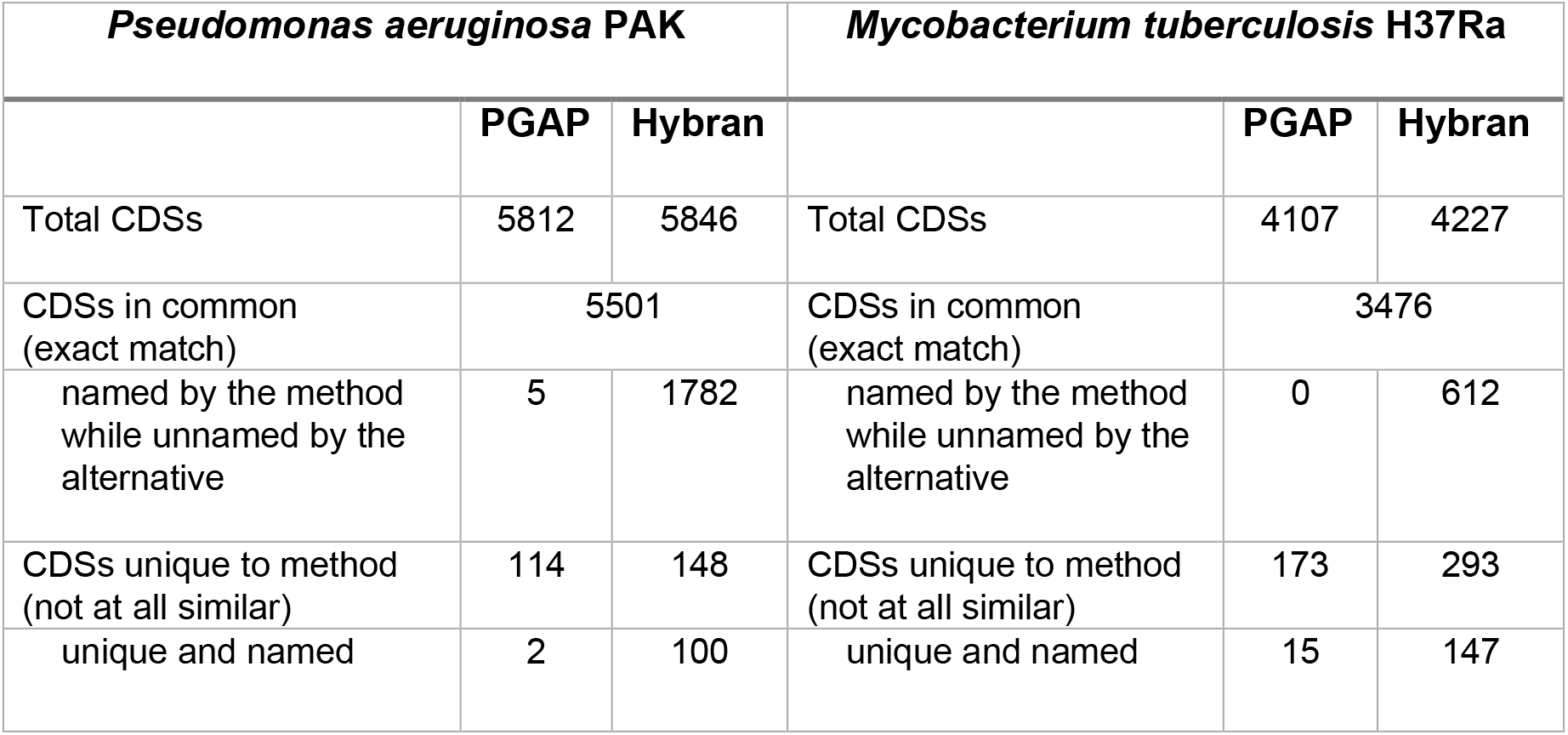
Results of PGAP and Hybran Annotation Comparison. Protein-coding sequence (CDS) annotations are compared for the two methods. Annotations that have been identically matched (by gene boundaries) are categorized as “in common”. CDSs that are “named” in PGAP are those with a non-empty gene name whereas, for Hybran annotations, they are those that have not been assigned a reference gene name rather than a generic (ORF#####) one. CDSs that are partially similar between each method are not compared here.

| *Pseudomonas aeruginosa* PAK |  |  | *Mycobacterium tuberculosis* H37Ra |  |  |
| --- | --- | --- | --- | --- | --- |
|  | PGAP | Hybran |  | PGAP | Hybran |
| Total CDSs | 5812 | 5846 | Total CDSs | 4107 | 4227 |
| CDSs in common (exact match) | 5501 |  | CDSs in common (exact match) | 3476 |  |
| named by the method while unnamed by the alternative | 5 | 1782 | named by the method while unnamed by the alternative | 0 | 612 |
| CDSs unique to method (not at all similar) | 114 | 148 | CDSs unique to method (not at all similar) | 173 | 293 |
| unique and named | 2 | 100 | unique and named | 15 | 147 |

For both genomes, Hybran also tended to find more CDSs uniquely and overall. For PAK, Hybran found 148 CDSs not in PGAP and assigned names to 100, over two-thirds, of them, whereas PGAP found 114 CDSs that Hybran did not, but assigned names to only 2 of them. For H37Ra, Hybran uniquely found nearly 300 CDSs and named approximately half of them (147/293), while PGAP named 15 out of its 173 unique calls. While uniquely called CDSs might at first seem to be candidate false positives, Hybran’s functional annotation of many of its unique calls suggests that many of them are not so.

## Discussion

Hybran is not a traditional genome annotation tool in that it does not prioritize functional annotation of genes. Its purpose, rather, is to facilitate pangenomic and evolutionary analyses. Even though a gene’s function may be unknown, the act of labeling this gene, by naming it accordingly, as orthologous to a hypothetical gene also seen in in the reference annotation makes the genome annotation more relevant for these applications. In this view, the observation that Hybran was able to name more genes than PGAP shows a difference in objective between these two tools, as the expectation from users of pure ab initio methods like PGAP would be that genes with unknown function should not be assigned a name.

Hybran is not fundamentally limited to Prokka as its source of ab initio calls. Future development can see Hybran extended to instead draw them from different tools such as Bakta(20), which has recently succeeded Prokka in its role as a rapid command line annotation tool, or even PGAP itself.

Liftoff is an alternative to RATT for reference-based annotation. RATT’s synteny-based approach has a great advantage for features difficult to independently align, but it will not capture, for example, a gene that has been duplicated or relocated because that change results in a break in synteny. Liftoff, on the other hand, may find such genes and all their occurrences, so long as they are not difficult to align. And in its searching for the reference genes across the sample’s entire genome, Liftoff is not limited to identifying genes at boundaries delineated by an ORF caller. Therefore, it represents a complementary approach both to RATT and to ab initio methods, so a future version of Hybran may incorporate it.

The objective of Hybran is to not only to produce a standard genome annotation, but one that captures and highlights relationships between strains emanated by experimental data hitherto underused.

## Funding

This work was funded by grants from National Institute of Allergy and Infectious Diseases (NIAID Grants R01AI105185 & R01AI163202). The funding bodies had no role in the design of the study or collection, analysis, and interpretation of data or in writing the manuscript.

## Acknowledgements

We would like to thank Theron James for code contributions and testing, and Derek Conkle-Gutierrez and M. Omar Din for thorough testing of the software.

## Notes

### Competing Interest Statement

The authors have declared no competing interest.

